# EEG frequency tagging reveals the integration of dissimilar observed actions

**DOI:** 10.1101/2024.05.10.593473

**Authors:** Silvia Formica, Anna Chaiken, Jan R. Wiersema, Emiel Cracco

## Abstract

Extensive research has demonstrated that visual and motor cortices can simultaneously represent multiple observed actions. This ability undoubtedly constitutes a crucial ingredient for the understanding of complex visual scenes involving different agents. However, it is still unclear how these distinct representations are integrated into coherent and meaningful percepts. In line with studies of perceptual binding, we hypothesized that similar movements would be more easily integrated. To test this hypothesis, we developed an EEG frequency tagging experiment in which two hand movements were displayed simultaneously at two different presentation rates. Crucially, the degree of similarity between the two movements varied along two dimensions, namely action identity (i.e., same or different performed movement), and agent identity (i.e., one agent performing a bimanual movement, or two agents moving each one hand). Contrary to our predictions, we found a larger intermodulation oscillatory component, indexing the integrated processing of the two individual movements, when they were less similar. We propose that integration-by-dissimilarity might serve as a top-down process to solve conflict caused by incongruent movements in a complex social scene.

## Introduction

During the vast majority of our daily activities, we are immersed in a social context: successfully navigating our environment inevitably requires quick and efficient interpretation of others’ actions. Correctly attributing thoughts, feelings, and intentions to our social partners is a complex and multifaceted process, involving the integration of several bodily features, such as facial expressions, gaze direction, posture, and limb orientation (Pitcher & Ungerleider, 2021). One crucial and rich aspect of social understanding is the processing of other’s movements (Pavlova, 2012). Literature supports the existence of specialized cognitive and neural mechanisms that are dedicated to processing biological motion (Blake & Shiffrar, 2007; Caspers et al., 2010; Giese & Poggio, 2003; Rizzolatti & Craighero, 2004). More recently, a growing body of studies is providing complementary and striking evidence that the brain not only processes isolated biological motion, but can also successfully represent the actions of distinct observed agents in visual and motor regions. This phenomenon has been indexed as an increase in imitative tendencies (Cracco et al., 2015, 2018; Cracco & Brass, 2018), and corticospinal excitability (Cracco et al., 2016) when observing two or more moving hands compared to one. Additionally, movement-specific patterns of activity have been shown in visual and motor regions while observing distinct hand postures, supporting the idea that different movements can be simultaneously represented by the brain (Cracco et al., 2019). These findings raise the intriguing question of how this discrete encoded information is organized in a coherent and meaningful way. Recent evidence shows that spatiotemporal relationships among multiple moving agents, and more specifically the synchrony of their movements, can be a driving factor in social grouping (Cracco, Lee, et al., 2022; Cracco, Papeo, et al., 2023). However, complex social scenes often include several agents performing different movements simultaneously, and understanding them must involve integrative processes that go beyond representing each individual observed movement.

Research in the field of early visual processing has firmly established that perceptual binding mechanisms are in place to parse the observed scene, tying together low-level features of objects, such as color and shape, in coherent perceptual units (Wagemans, Elder, et al., 2012; Wagemans, Feldman, et al., 2012). Analogous binding principles have been observed also at higher levels of the visual hierarchy. When parsing the elements of a complex visual scene, objects that are commonly used together or share a function can be perceptually grouped and processed as single entities, such as, for example, a pitcher pouring water into a glass (Green & Hummel, 2003, 2006; Kaiser et al., 2014; Kim & Biederman, 2011). These studies provided compelling evidence indicating that processing the relational properties among objects allows for a more efficient interpretation of the scene (Kaiser & Peelen, 2018).

In the last decade, EEG frequency tagging gained more popularity to investigate the phenomenon of perceptual binding. This technique is designed to capture the brain responses associated to rhythmically presented stimuli (Norcia et al., 2015; Vialatte et al., 2010). Notably, this approach can measure not only the oscillatory responses induced by different stimuli presented at different frequencies, but also neural signatures of their integration. These so-called intermodulation components emerge at combinations of the individual stimulation frequencies (e.g., their summation) and are taken as indication that the stimuli are being processed as an integrated entity (Boremanse et al., 2014), making this technique particularly apt to study perceptual binding. Across a broad range of visual stimuli (e.g., faces, illusory shapes, and moving dots), the emergence of intermodulation components has been taken as evidence that individual parts are processed as coherent wholes when they share specific visual properties, such as orientation and alignment (Alp et al., 2016; Boremanse et al., 2013, 2014; Palomares et al., 2012). For instance, Boremanse and colleagues (2014) showed reduced intermodulation components when observing two face halves belonging to distinct individuals flickering at different frequencies, as opposed to coherent faces formed by halves belonging to the same individual. These results demonstrated that the neural processing of coherent wholes is dissociated from the processing of its parts, and possibly reflects the emergence of an high-level representation (Boremanse et al., 2014).

The existence of perceptual binding mechanisms specifically dedicated to the processing of social scenes has been recently put forward (Papeo, 2020). It has been shown that the relational properties of individuals’ body postures (i.e., either facing towards or away from each other) play a role in inferring whether they are interacting (Papeo et al., 2017). Such binding of individuals into functional dyads leads to faster processing and enhanced memory of the social scene (Adibpour et al., 2021; Ding et al., 2017; Paparella & Papeo, 2022; Vestner et al., 2019, 2021) These results found initial support in EEG frequency tagging studies, revealing a larger intermodulation component when observing a dyad facing each other, as opposed to individuals facing away or inanimate objects (Adibpour et al., 2021; Goupil et al., 2023). However, non-social directional stimuli, such as arrows, can produce similar behavioral effects (Vestner et al., 2020, 2022). Therefore, whether this phenomenon arises simply as a consequence of bodies acting as directional cues (Vestner et al., 2022), or from binding mechanisms in place to process social interactions as integrated units, remains an object of investigation.

Crucially, this research focused exclusively on static body postures, and disregarded the vast informational value embedded in dynamic stimuli and the potential role of motion in making sense of complex social scenes (Pitcher & Ungerleider, 2021). To address this, frequency tagging has been fruitfully employed with moving social stimuli, such as point-light walkers (Alp et al., 2017; Cracco, Oomen, et al., 2022; Cracco, Papeo, et al., 2023; Troje, 2002), or sequences of rapidly alternating images of body postures to produce apparent movement (Cracco, Lee, et al., 2022; Cracco, Linthout, et al., 2023; Orgs & Haggard, 2011). Overall, these results suggest that the spatiotemporal properties of multiple moving social stimuli are used as perceptual cues, facilitating their grouping into units.

In the current study, we aimed at extending this body of research, and to test whether different sources of biological motion are integrated based on another well-known perceptual grouping principle, namely similarity (Wagemans, Elder, et al., 2012). We hypothesized that to efficiently handle multiple incoming streams of social information, the brain parses and integrates them into coherent units if they are similar and related. Therefore, we developed a frequency tagging paradigm in which identical or different hand movements, namely finger lifting or finger swiping, were presented simultaneously on the screen, moving at two different frequencies (F_1_ and F_2_). Additionally, the moving hands could belong either to the same or two clearly distinct actors. This setup allowed us to investigate brain responses associated to the two individual movements (stimulation frequencies F_1_ and F_2_), and their intermodulation components (F_1_ + F_2_ or other combinations of the two), assumed to reflect the binding of the two movements. We reasoned that if the brain groups and processes together moving stimuli based on the principle of similarity, more similar observed movements would result in a larger intermodulation response indicating their integrated processing. Accordingly, we expected the intermodulation component to track both our similarity manipulations, namely movement and agent identity. Two identical movements should share a higher degree of similarity compared to two different movements, thus being more readily integrated. Analogously, bimanual actions performed by a single agent are expected to be more similar than a dyadic movement, thus eliciting a larger intermodulation component.

## Methods

### Participants

A sample of 33 participants was initially recruited from the experimental pool of Ghent University. Eligibility criteria included no history of neurological or psychiatric disorder and normal or corrected-to-normal vision. All participants expressed their informed consent before the beginning of the experiment, and the whole protocol was approved by the ethics board of the Faculty of Psychological and Educational Sciences at Ghent University (2020/168). Sample size calculation was performed using the pwr package in R (Champely et al., 2020) and based on a medium effect size of dz = 0.50, aiming for a power of 80%. Due to poor EEG data quality (> 10% of the electrodes requiring interpolation), we discarded data from 3 participants, and one additional participant dropped out of the experiment before completing the task, thus resulting in a final sample of N = 29 (24 females, 5 males, M_Age_ = 22.48 ± 3.88, range_Age_ = 18 - 32). Participants received a monetary compensation of 25 € for their time.

### Task, stimuli, and procedure

The experiment was programmed using the Psychopy toolbox version 2021.2.3 (Peirce, 2007). Participants performed the task sitting comfortably in front of a computer monitor, and observed stimuli appearing on the screen. These consisted of rapidly alternating images of two hands, displayed simultaneously on a grey background to the left and the right of a grey midline running vertically. Each image was 512×767 pixels in size (Figure 1C). The rapid alternating between a sequence of two images with two different hand positions produced the illusion of movement. Two types of movements were presented, namely a swiping motion, consisting of a horizontal displacement of the index finger; and a tapping motion, showing the index finger being lifted vertically. Crucially, the two static hand images on the left and on the right side of the screen alternated at two different rates. On one side, images alternated every 24 frames (F1 = 2.5Hz, 400 ms) and on the other every 19 frames (F2 = 3.16Hz, ∼317 ms) (Figure 1B). These two frequencies were selected for several reasons. First, they aligned to the monitor 60 Hz refresh rate. Second, lower frequencies have been shown to produce larger and more reliable responses to high-level visual stimuli (Norcia et al., 2015). Finally, the first harmonics of the two frequencies did not overlap with each other, nor with the intermodulation component arising from their sum (*f1 + f2* = 5.66Hz), and were outside of the alpha range (8-12 Hz) to avoid confounds with attentional processes (Jensen & Mazaheri, 2010).

**Figure 1.**
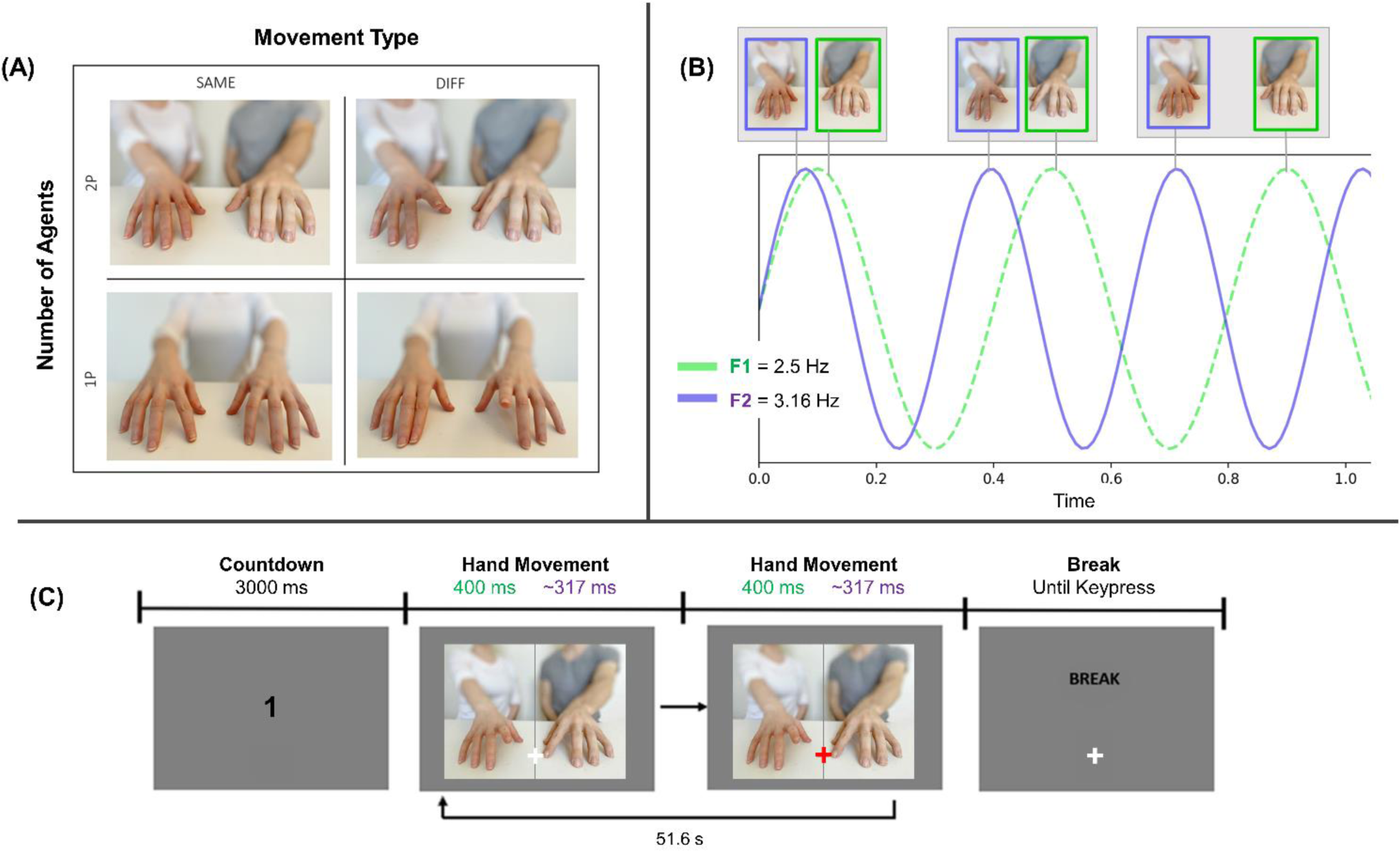
Experimental design. *Notes*. (A) 2 x 2 Factorial design with factors Movement type (*same* vs *different*) and Actor Number (*1P* vs *2P*). (B) Each of the two hands on the screen moved at a specific frequency, 2.5 Hz and 3.16 Hz, counterbalanced across blocks. (C) Timeline of an example trial. Each trial started with a countdown lasting 3 seconds, that was followed by images of the two hands, alternating at the specified frequencies to create the illusion of movement, for 51.6 seconds. The first and last three seconds consisted of fade in and fade out periods. Occasionally, the fixation cross turned red and prompted participants to press the spacebar. After each trial, participants were invited to have a self-paced break.

Each trial started with a 3-seconds countdown, during which the decreasing numbers appeared in the middle of the screen. This interval prompted participants to start focusing on the white fixation cross appearing along the midline of the screen, approximately in between the moving fingers. Next, the two hands moved for 51.6 seconds (i.e., 3096 frames), of which the first and last 3 seconds consisted of fade in/out periods in which image opacity was gradually decreased and increased, respectively. Participants were instructed to fixate on the cross, and to press the spacebar when it turned red. These changes occurred 5 to 8 times for each trial, and their duration was 400 ms. The goal of this color switch detection task was to ensure participants remained focused and paid attention to both moving fingers. Each trial was followed by a self-paced break, for a total duration of the whole experiment of approximately 32 minutes (Figure 1C).

The experiment consisted of a 2×2 factorial design. The first factor we manipulated was MOVEMENT TYPE, as the two hands could be performing the *same* movement (i.e., both tapping or both swiping) or two *different* movements (i.e., one tapping and one swiping). The second factor we manipulated was AGENTS, as the two moving hands could belong to one person (*1P*) performing the movements bimanually, or to different individuals (*2P*, Figure 1A), each performing one of the two movements. The two actors (one male, one female) were rendered clearly distinguishable based on their body posture and the color of their clothes. Each of the 4 experimental conditions resulting from our factorial design had 4 possible configurations, depending on which actor was displayed in condition 1P (actor 1 and actor 2), and the relative position of the actors in condition 2P (i.e., actor 1 on the right and actor 2 on the left and vice versa). The experimental session consisted of 2 blocks. In each of them all 16 combinations were presented in a randomized order. Therefore, the total number of trials for each participant was 32 (4 experimental conditions x 4 configurations x 2 blocks). One crucial difference between the blocks was the side of each frequency stimulation. If in the first block the hand on the left side moved at frequency F1 and the hand on the right at frequency F2, then in the second block this pattern reversed. The assignment of frequency stimulation was counterbalanced across participants.

### EEG Recordings and Preprocessing

Electrophysiological data were recorded with Ag/AgCl active electrodes, using an ActiCHamp amplifier and BrainVisionRecorder software (version 1.21.0402, Brain Products, Gilching, Germany), at a sampling rate of 1000 Hz. The placement of electrodes was consistent with the 10-20 international system (Klem et al., 1999), with the exception that electrodes TP9 and TP10 were replaced by OI1h and OI2h to ensure better coverage of posterior scalp sites. Two additional external electrodes were placed above and below the left eye to record vertical eye movements. Fpz was used as ground electrode, while Fz served as online reference.

Offline processing was performed with the Letswave 6 toolbox (https://www.letswave.org/). First, a bandpass fourth-order Butterworth filter was applied to the raw data, retaining frequencies within 0.1 Hz and 100 Hz. Data was then segmented time-locked to the onset of the hand motion, including 2 seconds before and after the video (−2 to 53.6 seconds). An independent component analysis (ICA) was performed on the merged segmented data to remove electrophysiological activity associated with horizontal and vertical eye movements. On average, 1.76 (±0.68) components were removed for each participant (range: 1 - 3). Following ICA, data were visually inspected to detect noisy electrodes, that were then interpolated using the average of the three nearest neighboring channels. An average of 1.96 (±2.07) electrodes were interpolated across participants (range: 0 - 6). Afterwards, electrode Fz that was used as online reference was reinserted for all participants and the data was re-referenced to the average of all 64 electrodes. Finally, the segments were cropped to exclude the ramp in and ramp out periods, resulting in epochs of 45.6 seconds (3 to 48.6 seconds following movement initiation). This duration ensured that epoch length was a multiple of both base frequencies, so that it contained an exact number of cycles.

To investigate the base motion frequencies (i.e., 2.5 Hz and 3.16 Hz) and their harmonics, epochs were averaged in the time domain within each block. These time-series were then transformed into the frequency domain by applying a discrete Fourier Transform (FFT), and the amplitudes (μV) were normalized (i.e., divided by N/2, where N is the length of the data). Averaging across blocks was then performed in the frequency domain, after adjusting electrodes labels so that they reflected brain responses ipsilateral and contralateral to the stimulated hemifield, rather than right and left scalp locations. This procedure was motivated by the strong lateralization of the base motion frequencies responses based on which visual hemifield was being stimulated, and ensured we did not cancel out stimulation-related activity due to the counterbalanced presentation side of the two frequencies.

On the contrary, the IM response did not exhibit any lateralization. Therefore, we averaged trials per condition across both blocks in the time domain. We then transformed the time-series into the frequency domain, applying an FFT with normalized (N/2) amplitudes. This procedure also allowed us to average together in the time domain more trials with respect to responses to the base frequencies. It has been previously shown that averaging together large sets of trials is beneficial to detect more robustly IM components, which can be orders of magnitude smaller than the base frequencies (Boremanse et al., 2013).

Finally, the frequency spectra obtained with this procedure were noise corrected by implementing a signal-to-noise subtraction (SNS). We subtracted from the amplitude at each frequency bin the average amplitude of the twenty surrounding bins (10 per side, excluding the immediately adjacent ones). The SNS method attenuates low frequencies artifacts and results in more defined spectral peaks (Norcia et al., 2015). The resolution of the amplitude spectra and corresponding SNS is equal to the FFT bin size, defined as the inverse of the epoch length, and hence was ∼0.02 Hz (1/45.6).

### Frequencies selection

Entraining a specific frequency by means of frequency tagging elicits a neural response not only at said frequency (F), but also at its harmonics (2F, 3F, …). It has been shown that the best approach to capture the brain response to a tagged frequency is to sum the baseline-subtracted amplitudes of all relevant harmonics (Norcia et al., 2015; Retter et al., 2021; Retter & Rossion, 2016). To determine which harmonics to include for each tagged frequency and their intermodulation component, we implemented a data driven procedure to capture periodic responses significantly above noise levels (Cracco, Lee, et al., 2022; Cracco, Oomen, et al., 2022; Retter & Rossion, 2016). First, we computed the grand-averaged amplitude across all participants, conditions, and electrodes. Each frequency bin was then z-scored using the 20 surrounding bins (10 for each side), excluding the immediately adjacent ones. Finally, the peaks in the amplitude spectrum were retained if they had a z-score > 2.32, corresponding to p < 0.01. With this procedure, we identified 8 frequencies for each of the two base frequencies 2.5 Hz and 3.16 Hz (Figure 2, left panel). Subsequent analyses concerning base frequencies were carried out on the summed amplitudes of these harmonics.

**Figure 2.**
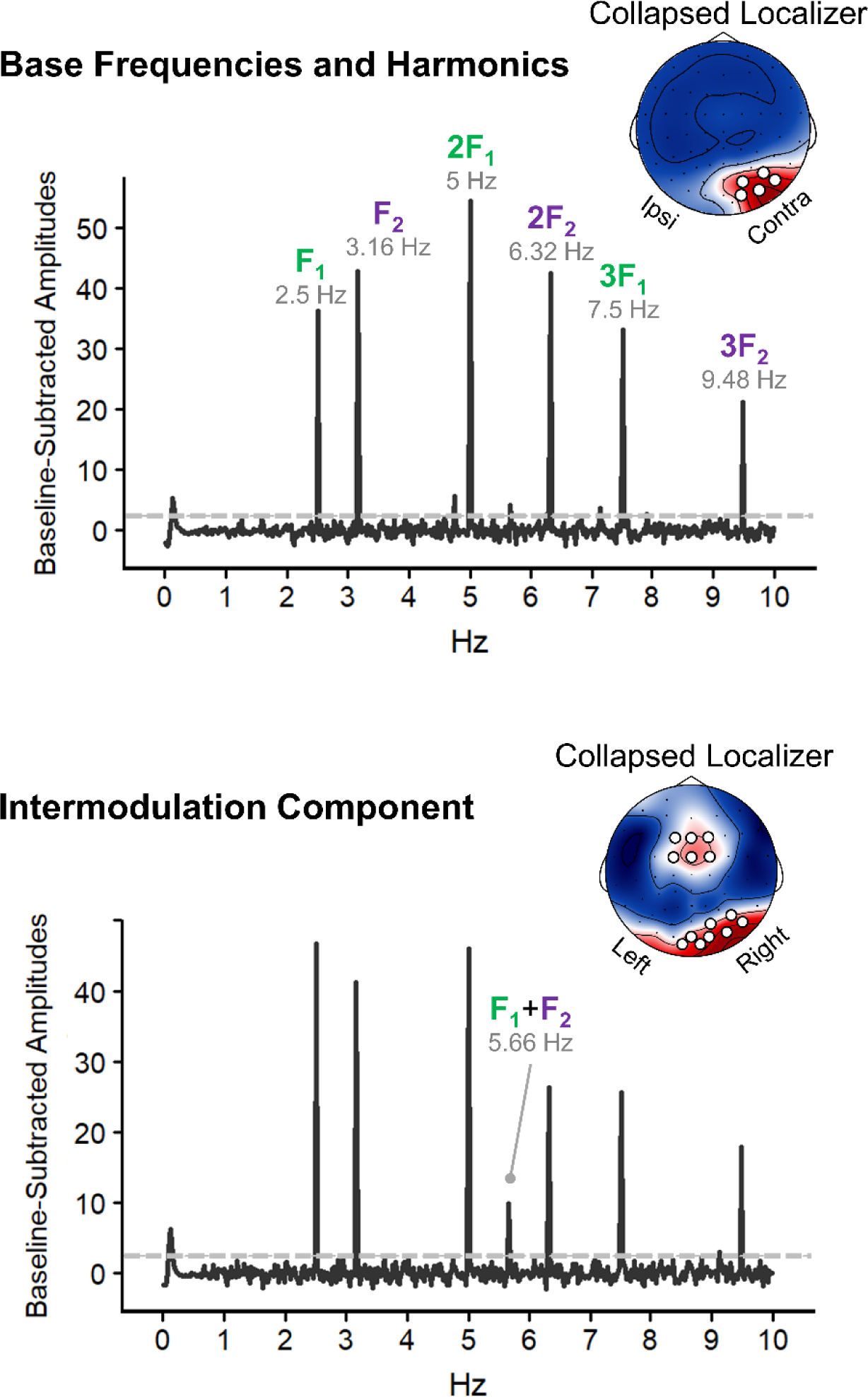
Frequencies and Electrodes Localizers. *Note.* Each panel shows the z-scored Baseline-subtracted amplitudes, plotted for the electrodes resulting from the collapsed localizers. Results for the collapsed localizers are displayed in the inset topographies, maximal amplitudes are in red. Larger white dots indicate the selected electrodes. The gray horizontal line indicates the significance level of 2.32 above which harmonics are retained for subsequent analyses. *Upper panel:* Results of frequencies and electrodes selection for the two base frequencies. Only the first three harmonics of each base frequency are displayed. *Lower panel*: Results of frequencies and electrodes selection for the intermodulation component. The only retained frequency is 5.66 Hz, resulting from the summation of the two base frequencies.

To identify relevant frequencies for the intermodulation component we focused on the first three harmonics of F_1_ and F_2_ combinations. We then adopted the same z-scoring procedure outlined above, and retained only the intermodulation frequency 5.66 Hz (F_1_ + F_2_, Figure 2, right panel).

### Electrodes selection

To select a subset of relevant electrodes for our analyses, we implemented a collapsed localizer approach (Cracco, Lee, et al., 2022; Cracco, Oomen, et al., 2022; Luck & Gaspelin, 2017). The goal of this approach is to average the scalp topographies of the summed baseline-corrected amplitudes across all participants and conditions, and then pick the electrodes showing the most marked activations. After seeing that the two base frequencies (i.e., 2.5 Hz and 3.16 Hz) had nearly identical scalp topographies, their amplitudes were averaged together. Notably, their responses appeared to be lateralized and prominent in the occipital electrodes contralateral to the stimulated hemifield. Therefore, we flipped the electrode coordinates for blocks in which the frequency being analyzed was presented in the right hemifield. In this way, electrodes labeled as belonging to the right hemisphere displayed responses contralateral to the stimulation location, and, conversely, left scalp electrodes reflected ipsilateral responses. We identified five electrodes showing a strong response, namely PO4, PO8, P6, P8, and O2, and conducted all analyses concerning base frequencies on the averaged amplitudes across them (Figure 2, left panel inset).

With respect to the intermodulation component, we adopted the same collapsed localizer approach (without flipping electrodes label, as responses were not lateralized), and detected two regions of interest. The first one was occipital, including electrodes PO4, PO8, O2, Oz, OI2H, OI1H, P6, and P8; while the second was observable over midfrontal electrodes FCz, FC1, FC2, F1, Fz, and F2 (Figure 2, right panel inset). Within each of these two regions, the amplitudes were averaged across the included electrodes.

### Data analysis

All statistical analyses were carried out with R (version 4.1.2). To investigate the brain response to the two base frequencies (and their harmonics), we conducted a repeated measures ANOVA with Frequency (2.5 Hz vs 3.16 Hz), Movement Type (same vs different), and number of Agents (1P vs 2P) as within-subject factors.

Regarding the intermodulation component, the baseline-subtracted amplitudes of frequency 5.66 Hz were submitted to a repeated measures ANOVA with factors Region (frontal vs occipital clusters), Movement Type (same vs different), and number of Agents (1P vs 2P) as within-subject factors.

## Results

### Behavioral Results

Participants were asked to fixate on the cross displayed along the midline, and to monitor its color to detect changes throughout the whole duration of the trial. Upon change detection, they had to press the spacebar. Participants had high detection accuracy, reaching on average 98.75% (±1.93). On average, participants pressed the spacebar to indicate a detected color change in absence of an actual change (i.e., false alarms) 1.06 times (±0.23) throughout the whole experiment.

### Base Frequencies 2.5 Hz and 3.16 Hz

Amplitudes at the base frequencies and their harmonics reflect brain activity in response to the individual hand motions. The 2 (Frequency: 2.5 Hz vs 3.16 Hz) x 2 (Movement Type: same vs different) x 2 (Number of agents: 1P vs 2P) repeated measures ANOVA revealed only a significant main effect of Number of agents, F_1, 28_ = 8.77, p = 0.006, d_z_ = 0.55, indicating higher amplitudes for trials with two distinct agents. All other main effects and all interactions were not significant (Fs < 2.36 and ps > 0.13). Since we did not observe a main effect of Frequency, we collapsed across the two Frequencies in Figure 3.

**Figure 3.**
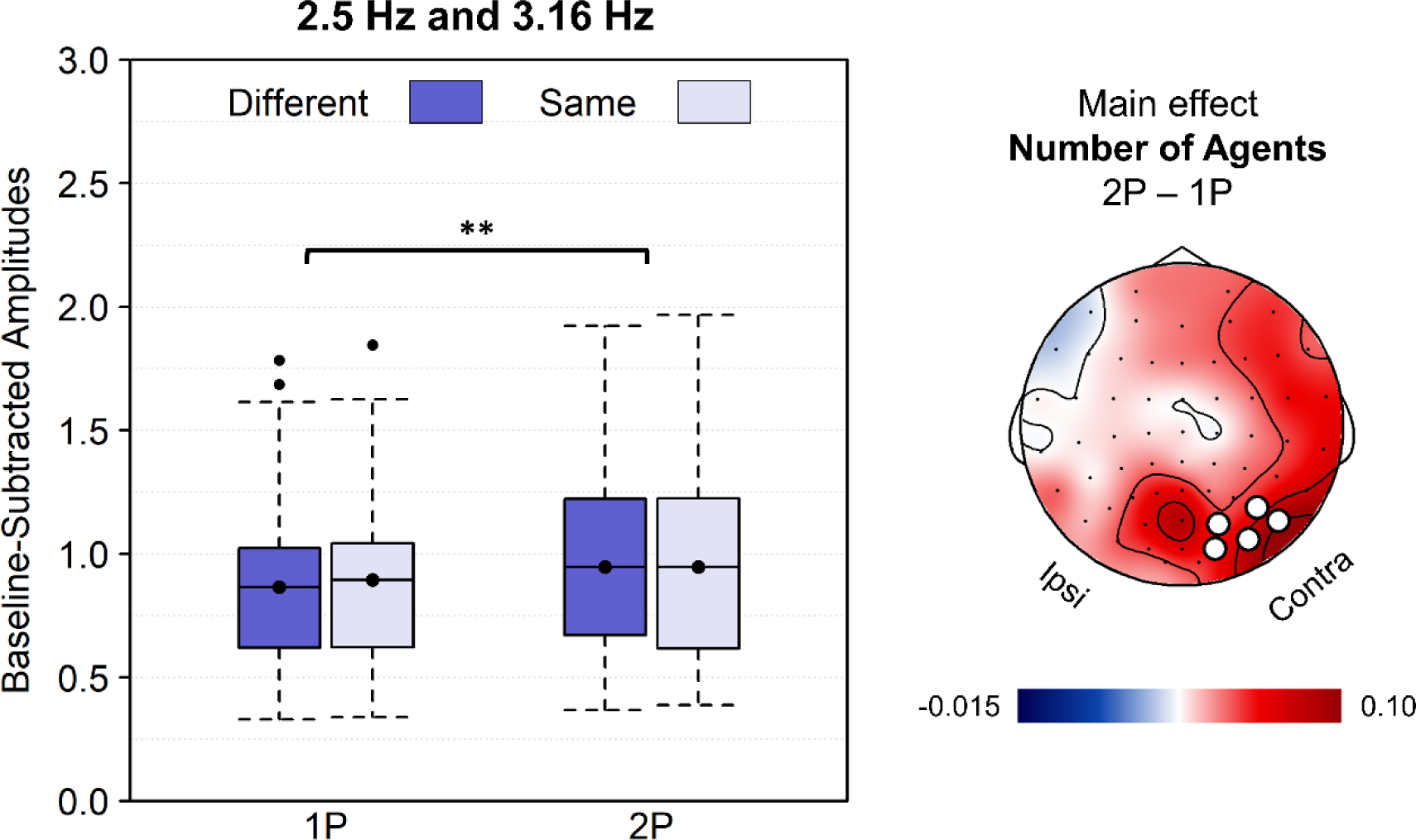
Base Frequencies Results. *Note*. The left panel depicts the baseline-subtracted amplitudes, averaged across the two base frequencies (2.5 Hz and 3.16 Hz) and their harmonics, for the cluster of electrodes PO4, PO8, P6, P8, and O2. The asterisks indicate the main effect of Number of agents (p = 0.006). The right panel shows the topography of the neural response for the main effect of Number of agents, obtained by subtracting the response to trials with one agent from trials with two agents. Note that the electrodes locations have been manipulated in order to collapse across lateralized activity, and display contralateral activity on the right hemisphere and ipsilateral activity on the left hemisphere. The electrodes highlighted in white are those obtained from the collapsed localizer and used for the analyses.

### Intermodulation component 5.66 Hz

Neural response to frequency F_1_ + F_2_ (i.e., 5.66 Hz) was characterized by two main foci of activity, one more parietal-occipital and one over midfrontal electrodes. Both were included in a repeated measures ANOVA with a factor Region (occipital vs frontal), alongside factors Movement Type (same vs different), and Number of agents (1P vs 2P). We observed a main effect of Movement Type (F_1, 28_ = 6.20, p = 0.019, d_z_ = 0.46), indicating a larger response to different compared to identical movements. Moreover, the effect of Number of agents was also significant (F_1, 28_ = 6.88, p = 0.014, d_z_ = 0.49), reflecting an enhanced response when observing two distinct actors compared to one actor performing a bimanual action. All other effects did not reach significance (Fs < 2.52, ps > 0.12, Figure 4).

**Figure 4.**
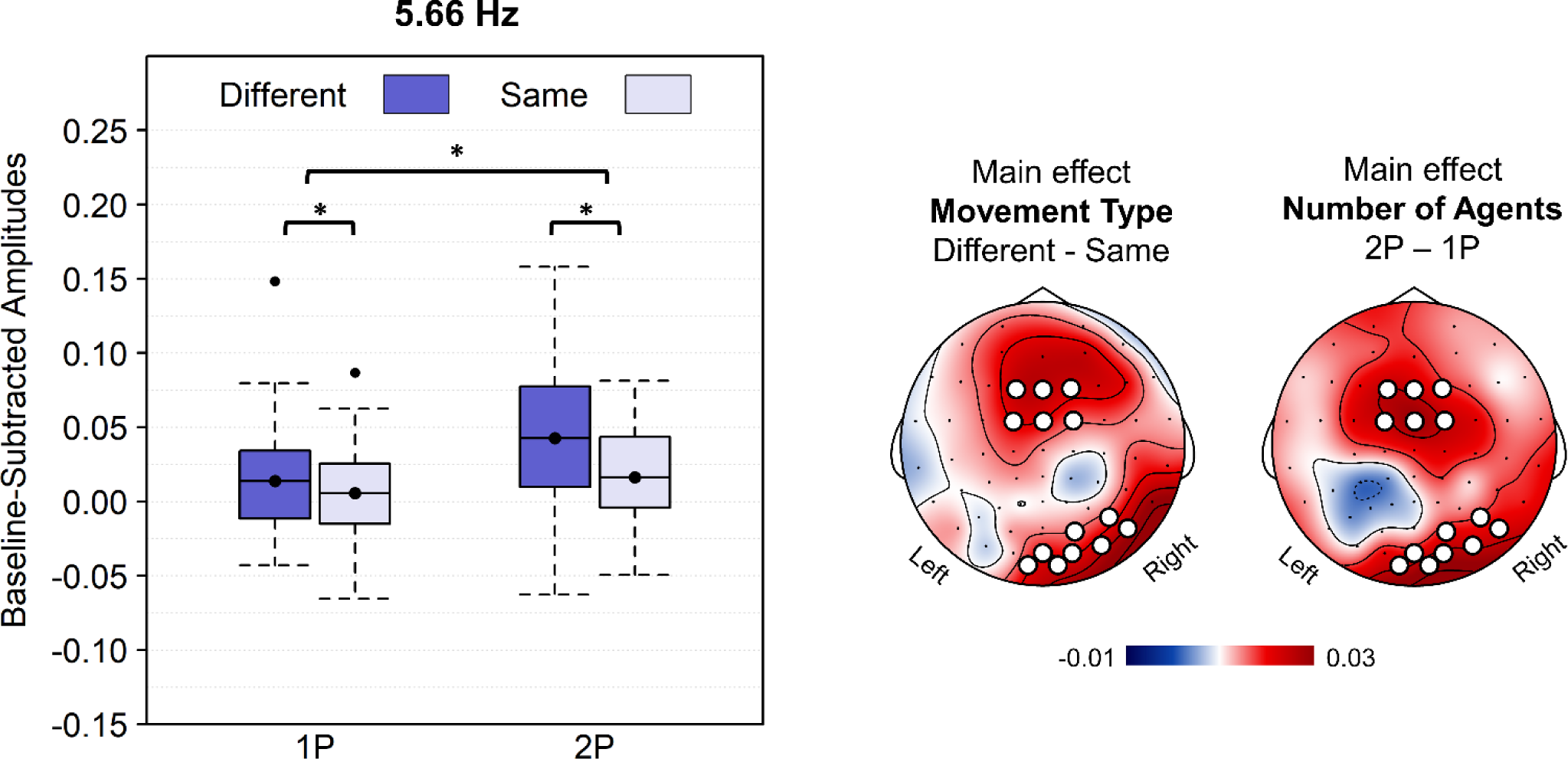
Intermodulation Component Results. *Note*. The left panel depicts the baseline-subtracted amplitudes, averaged across the two considered clusters of electrodes (occipital and frontal), for the intermodulation frequency 5.66 Hz. The asterisks indicate the main effect of Number of agents (p = 0.014), and Movement Type (p = 0.019). The right panel shows the topographies of the neural response for the two main effects. The electrodes highlighted in white are those obtained from the collapsed localizer and used for the analyses.

## Discussion

A growing body of research consistently indicates that multiple observed movements can be simultaneously represented in the brain (Cracco et al., 2015, 2016, 2019; Cracco & Brass, 2018). However, how such complex incoming information is parsed and integrated into meaningful perceptual entities is still unclear. Recent evidence is suggesting that the same binding principles that have been widely described in the field of visual perception (Wagemans, Elder, et al., 2012; Wagemans, Feldman, et al., 2012) might be at play also when processing complex social stimuli (Cracco, Lee, et al., 2022; Cracco, Papeo, et al., 2023). In the present study, we investigated whether the principle of similarity plays a role in making sense of multiple observed movements. More specifically, we hypothesized that identical movements are more likely to be processed in an integrated way, especially when they are executed bimanually by the same agent. On the contrary, different movements performed by distinct agents were expected to share the lowest level of similarity, thus inducing less integration processes. To test this hypothesis, we developed a frequency tagging paradigm, in which two hands moved simultaneously on the screen at two different frequencies. Crucially, the two hands could be executing the same or two different movements, and could belong to one agent or two agents. With this experimental design we set out to dissociate the neural responses to each individual movement and to their integration. The two similarity manipulations we implemented (same vs different movements and 1 vs 2 agents) tackled two slightly different questions. While the former was aimed at testing directly if movements identity has a role in their integration, the latter investigated whether integratory processes are at play to interpret social scenes involving multiple agents.

We found that the intermodulation response at 5.66 Hz, reflecting the processes integrating the two individual moving stimuli (i.e., F_1_ + F_2_), was significantly influenced by both our manipulations. However, in stark contrast to our predictions, this response was larger when observing two different movements compared to the same movement, and when these were performed by two distinct agents rather than by the same agent bimanually. Although these results confirm the recruitment of integrative processes when observing multiple movements, they diverge from the typical findings in the perceptual binding literature, which consistently showed increased IM responses under conditions of similarity and alignment (Alp et al., 2016; Boremanse et al., 2013, 2014). Conversely, in the present study it appears to be dissimilarity, rather than similarity, to trigger and drive the integration of observed actions.

One possible explanation for such a reversed effect is that congruent pairs of movements are represented in the brain as a single movement, thus not requiring integration. Rather than encoding twice the same action, the brain is likely adopting the more efficient strategy of representing the movement only once, while relying on univariate shifts of this activation pattern to index the number of observed movements. In line with this perspective, previous studies on motor resonance revealed larger activation of the primary motor cortex, quantified by motor-evoked potentials, when observing two hands performing the same movement compared to a single moving hand (Cracco et al., 2016). This hypothesized representational mechanism is also consistent with findings indicating a plateau in the facilitating effect of observing multiple congruent movements (Cracco, Bernardet, et al., 2022; Cracco & Brass, 2018). The increase in activation level can follow a linear trend only up to the motor threshold, as its trespassing would result in the execution of the same motor plan. Consequently, differences in motor activation and ensuing facilitating effects are larger between small numbers of observed agents, and reach a plateau for bigger groups. On the contrary, when observing different movements, these need to be separately represented in the brain. This would then require integrative processes to be in place to extract a coherent percept from the inconsistent incoming visual inputs.

Analogously, our data revealed also a more prominent IM response when observing two distinct agents, compared to one agent acting bimanually. It is reasonable that a single agent performing a bimanual movement is already represented in the brain as a unitary item. Conversely, integration processes would necessarily be recruited to a further extent when observing two separate agents acting alongside, as these would not be perceived as a single unit, but rather need to be combined into a moving dyad. From this perspective, such integration would be a core mechanism for social cognition, allowing the processing of multiple individuals as a group, if they share perceptual or functional properties.

Differences in the neural activity related to the number of distinct moving agents were noticeable also in response to the base stimulation frequencies. Again, a larger response in scalp location contralateral to the stimulation side was present when observing two compared to one agent. We think this result is likely reflecting a higher deployment of attentional resources to each of the two lateralized agents in the 2P condition. In fact, previous findings highlighted a more pronounced response to the flickering frequency of visual stimuli when the observer directed attention to them (Kashiwase et al., 2012; Zhigalov et al., 2019). This interpretation is consistent with the two agents being perceived as two separate entities, and thus being separately attended.

Altogether, our findings hint at the unpredicted idea that social binding mechanisms are triggered by incongruent incoming information. Rather than enhanced processing of social scenes that are already consistent (e.g., one agent performing congruent bimanual movements), we revealed integrative processes are needed to make sense of incongruent movements. These results indicate that conflict, rather than the lack thereof, could be one factor signaling the need for proactive integration. Interestingly, neuroimaging studies have shown that observing incongruent movements elicits activation of the anterior cingulate cortex (ACC), an area that has been broadly associated with conflict processing and with the exertion of cognitive control (Botvinick et al., 2001; Cracco et al., 2019). Consistently, the intermodulation component showed a frontocentral scalp distribution analogous to that observed in electrophysiological studies targeting conflict (Cavanagh & Frank, 2014). We speculate that the perceived motor conflict caused by observing two incongruent movements may trigger control processes aimed at integrating the discording actions into a coherent perceptual unit. Previous studies described the integrated processing of similar items as an emergent property of visual perception (Alp et al., 2016). Here, we put forward the proposal that in specific contexts also dissimilarity can be a relational property of visual stimuli that induce their integration. In these cases, integration can be seen also as a proactive mechanism exerted in a top-down fashion, as a tool for conflict resolution in complex social scenes.

It is crucial to reiterate that the results of the present study deviate from our original hypotheses. Although surprising, we interpreted them in light of related findings, and we think they pave the way for new exciting research questions. For example, whether the integration of dissimilar items is specific to dynamic social stimuli warrants further research. Moreover, which characteristics of the movements and the agents induce integrative processes should be pinpointed in more detail. In our study, we used meaningless hand motions, and the agents were shown only partially. This setup is clearly a simplification of the real-life movement dynamics we commonly engage with. More complex, full-body movements should be investigated to probe the role of different social features in this process. For example, it remains an open question to what extent movements that are incongruent but functionally related are integrated. (Sebanz et al., 2006). The integration-by-dissimilarity that we show in our current study constitutes a potentially crucial mechanism for the interpretation of complex social dynamics, allowing the integration of distinct agents into functional and meaningful units.

## Acknowledgements

SF was supported by the German Research Foundation (Deutsche Forschungsgemeinschaft, DFG) under Germany’s Excellence Strategy-EXC 2002/1, Science of Intelligence (Project Ref.: 390523135) and the Einstein Foundation Berlin. EC was supported by a senior postdoctoral fellowship (12U0322N) and a research grant (1500720N) awarded by the FWO.

